# Bacterioplankton diversity and pollution levels in the estuarine regions of Chaliyar and Anjarakkandi rivers, Kerala, India

**DOI:** 10.1101/2021.04.06.438743

**Authors:** D. Nikhitha, Arunaksharan Narayanankutty, Jisha Jacob

**Author notes:** **Corresponding author: Dr. Jisha Jacob, Ph.D**, Assistant Professor, Molecular Microbial Ecology Lab, PG and Research Department of Zoology, St. Joseph’s College (Autonomous), Devagiri, Calicut, Kerala, India.

## Abstract

Rivers and the estuarine regions play crucial roles in the environment by controlling the nutrient cycling, waste disposal, and also in supporting biodiversity. However, the estuarine regions of rivers are highly susceptible to disruptive changes by anthropogenic pollutants and eutrophication. Bacterioplankton diversity is the best indicators of the pollution level and eutrophication status of the rivers. The present study evaluated the eutrophication status and bacterioplankton diversity of the estuarine regions of Chaliyar and the Anjarakkandi originated from the Western Ghats. The present study observed the presence of increased incidence of bacterioplankton comprising of proteobacteria, actinobacteria and cyanobacteria in the estuarine regions of Anjarakkandi river compared to that of Chaliyar river. Further, the percentage incidence of faecal bacteria including Bacillus subtilis and E. coli, were also found to be in high in the Anjarakkandi river; in corroborating with this, the diversity of nitrogen metabolizing bacteria was also higher in the estuarine samples of Anjarakkandi river. Corroborating with this, the levels of water nutrients including nirates, nitrites and phosphates were high in the Anjarakkandi compared to the Chaliyar river. Together, it is possible that there might be increased faecal contamination and subsequent increased eutrophication in the estuarine regions of Anjarakkandi river. Altogether, the study concludes that the Anjarakkandi river may be more polluted and which is partly contributed by faecal contaminants and also responsible for the change in the bacterioplankton community of the estuaries.

## 1. Introduction

Indian sub-continent has a wide network of aquatic systems including small streams, rivers, lakes, marine waters and estuaries (Khadse et al. 2012; Khadse et al. 2010). Among these, several important rivers are being originated from the Western Ghats, which are important water resources in southern India (Ramachandra et al. 2018). However, the pollution levels of these south Indian rivers are increasing every year and which is most affected in their esturine regions (Gopal et al. 2018). Estuaries are one among the most urbanised ecosystem and greatly facing environmental pressure by anthropogenic activities includes habitat loss, algal blooms, anoxia, and contamination by sewage, heavy metals and other organic and inorganic pollutants (Bashir et al. 2020). They are also considered as the highly productive and diverse ecosystem, and possess unique microbial stress indicators which are vulnerable to anthropogenic contaminants (Duarte et al. 2017; Julian and Osborne 2018). Darmadam and Chaliyum are estuarine regions of Anjarakkandy and Chaliyar river respectively which are flowing through both rural and urban banks constitute anthropogenic contamination.

In usual scenario shows a ubiquitous distribution of bacterioplankton around the globe because of their small size, great abundance and easy dispersal. But some marine bacterioplankton possess geographical distribution pattern like macroscopic animals possess (Liu et al. 2015). Evaluating the distribution pattern of both particle associated and free living bacterioplankton are of higher significance, as the functioning of these bacteria remain key determinants of nutrient concentrations and fluxes by fixing carbon and nitrogen, assimilating and remineralizing dissolved and particulate organic matter in the ecosystem (Gómez-Consarnau et al. 2019). Hence, the diversity of bacterioplankton present in the aquatic environment can be an indicative of the quality of water and level of pollution in the water-body (Salloto et al. 2012; Zhao et al. 2020). Several methods are being utilized for the analysis of bacterioplankton; among these the predominant are culture independent methods (Li et al. 2019).

In recent years, microbial ecologists have effective ways to characterise microbial communities have been created. Numerous studies have indicated taxonomic variability in both time and space in the genetic diversity of bacterioplankton communities. However, because this data is focused largely on 16S rRNA gene sequences, the physiological characteristics and ecological functions of the genotypes identified remain largely unknown. The use of high performance sequencing technologies has made way for the study of the microbial group structure using amplicon sequencing and metagenomics. Hence, the present evaluated the bacterioplankton diversity in the estuarine regions of two selected rivers of Kerala employing high throughput pyro-sequencing techniques of 16s rRNA gene for culture independent bacterioplankton.

## 2. Materials and Methods

### 2.1 Study area and Sampling

Chaliyum and Darmadam are Esturine regions of Chaliyar and Anjarakkandi river respectively. Water samples were obtained from three stations of each site. Samples (post monsoon) were collected from 0.5m below the air -water interface using depth sampler. The collected samples were stored in a dark container and kept in 4°c immediately after the collection.Physico chemical parameters like Temperature (T), pH, Sainity, Total Dissolved Solids(TDS), Conductivity were measured using water analyser (EUTECH PC 450)and Dissolved oxygen(DO),Biochemical oxygen demand(BOD),Chemical oxygen demand(COD), of samplesof each stations were measured using standard methods(APHA 2005).

The samples from there stations were pooled and filteredthrough 3.0-μm-pore-size and 0.2 μm-pore-size membraneFiltersto collect particle-associated (PA) and free-living (FL) bacterioplankton, respectively and stored in - 20°c deep freezerafter filtration.The remaining sample of water from each stations of two sites were used for nutrient analysis. The filters were subjected to DNA extraction

### 2.2 Nutrient Analysis

Both organic and inorganic nutrients were measured by following the standard methods for the examination of water and waste water.Total hardness, Calcium, Magnesium were measured byEDTA titration method.Chloride(Argentometric method), Iron(Phenanthroline method), Nitrate (Phenol-disulphonic acid method), Sulphate(Turbidity method), Ammonium (Nesseler’s method),Phosphate(Stannous chloride method), Particulate organic carbon (POC) and Dissolved organic carbon(DOC) were found using Ash method (APHA 2005).

### 2.3 DNA Extraction and PCR amplification

DNA isolation from frozen filter were done using the EXpure Microbial DNA isolation kit (BogarBio Bee stores Pvt Ltd) by following manufactures instructions. We used lysis buffer to homogenise the cell.After the extraction DNA concentrations were measured by Qubit 3.0 and the DNA samples were stored at −20°C until further use.

16S rRNA genes was PCR amplified.from extracted DNAs using primers that were designed specifically for 454 high throughput pyrosequencing. The forward primer were 27F(5’AGAGTTTGATCMTGGCTCAG3’) and the reverse primer were 1492R(5’AAGGAGGTGATCCAGCCGCA3’). Composition of the Taq Master Mix consist ofTaq DNA polymerase is supplied in 2X Taq buffer 0.4mM dNTPs, 3.2mM MgCl2 and 0.02% bromophenol blue. A 25 μL reaction solution contains 5 μL of isolated DNA 1.5μL of Forward Primer and Reverse Primer, 5 μL of deionized water, and 12 μL of Taq Master Mix This PCR amplification consist of 25 cycles with initial 2 minutes denaturation(95ºc) and final 10 minute extension (72ºc). Each cycle had Denaturation (95°C 30 sec) Annealing (60°C 30 sec) Extension(72°C2min). Unincorporated PCR primers and dNTPs from PCR products removed by using Montage PCR Clean up kit. The quality, quantity (App.100ng/μl) and formulation of the PCR product was checked using Qubit Fluorometer 3.0.

### 2.4 Sequencing annotation and library coverage calculation

The pyrotag sequences obtained for each of the sample were sorted and low quality sequences were removed as per the previously described protocols. Further processing and taxonomic identification upto species level was done using BLASTn program based on the 16S rRNA sequence similarity as per the protocols described by Mou et al. (2013).

### 2.5 Statistical analysis

The values were expressed as mean± SD of eight different sampling sites from both the estuarine regions. The statistical analysis used paired t-test for comparison of nutrient quality of water samples from both the stations and a variation of p<0.05 is considered statistically significant.

## 3. Results

### 3.1 Nutrient composition of the estuaries

Changes in the various physical and chemical parameter along with the biological aspects of water quality from the estuarine regions of Anjarakkandi and Chaliyar river are listed in the Table 1. As shown in the table, there was limited variation in the temperature of water present in both the estuarine regions; however, the Anjarakkandi river had significantly reduced levels of dissolved oxygen content. Concomitantly, there observed an increased level of nitrate, ammonium, phosphate and sulphate. Further, there observed significantly elevated BOD and COD values in these samples (Table 1).

**Table 1.** The nutrient composition of the estuarine regions of Chaliyar and Anjarakkandi river, Kerala, India.

| Parameter | Chaliyar | Anjarakkandi |
| --- | --- | --- |
| Temperature(°c) | 31.0± 0.0 | 31.0± 0.00 |
| pH | 7.57 ± 0.02 | 7.72± 0.02* |
| Salinity (ppt) | 10.60 ± 0.00 | 13.67± 0.06*** |
| TDS (g/l) | 9.94 ± 0.00 | 12.63± 0.06*** |
| Conductivity (ms/cm) | 13.95 ± 0.06 | 22.06± 0.07** |
| Total Hardness (mg/ml) | 1548.33 ± 2.89 | 2150.33±0.058*** |
| DO (mg/l) | 6.35 ± 0.03 | 3.82± 0.04** |
| BOD (mg/l) | 2.07 ± 0.01 | 8.87± 0.21** |
| COD (mg/l) | 39.10 ± 0.17 | 77.83± 0.81*** |
| Calcium (mg/l) | 100.00 ± 0.00 | 158.33± 1.53* |
| Magnesium (mg/l) | 320.80 ± 3.82 | 426.00± 1.73** |
| Chloride (mg/l) | 5998.44 ± 0.49 | 8463.41± 0.13*** |
| Iron (ppm) | 1.42 ± 0.10 | 0.50± 0.00* |
| Nitrate (ppm) | 0.29 ± 0.03 | 0.46 ± 0.00* |
| Ammonium (ppm) | 0.31 ± 0.01 | 0.43± 0.00* |
| Sulfate (ppm) | 139.62 ± 0.06 | 186.62± 0.04* |
| Phosphate(ppm) | 0.03 ± 0.01 | 0.02± 0.00 |
| POC (%) | 0.21 ± 0.02 | 2.12± 0.00*** |
| DOC (%) | 0.14 ± 0.01 | 0.39± 0.01** |

### 3.2 Bacterioplankton diversity of the estuaries

The diversity of bacterioplankton in the samples is given in table 2 and 3. The kingdom level analysis indicated the presence of both particle-associated and free living bacterioplanktons. In the estuarine regions of Anjarakkandi river the presence of bacterial diversity was significantly higher than those present in the estuarine regions of Chaliyar river (Table 1). Further, the phylum level analysis indicated the diversity of proteobacteria (67.07%) and firmicutes (17.24%) in the estuaries of Anjarakkandi river. On contrary, the diversity of proteobacteria and firmicutes in the Chaliyar river range was 46.35 and 31.51%, which was significantly lower than that of the Anjarakkandi river. In the species level identification, the faecal bacteria Bacillus subtilis was much higher in the Anjarakkandi river estuaries with 16.19%; the same in Chaliyar river was 10.07% (Supplementary material 1).

**Table 2.** Changes in the diversity of particle-associated and free living bacterioplankton in the estuarine regions of Anjarakkandi and Chaliyar river of Kerala, India

|  | Kingdom level |  |  | Phylum level |  |  |
| --- | --- | --- | --- | --- | --- | --- |
|  | Bacteria | Archaea | Eukaryota | Proteobacteria | Firmicutes | Euryarchaeota |
| Anjarakkandi- PA | 1768 | 217 | 157 | 1067 | 572 | 181 |
| Anjarakkandi- FL | 1027 | 78 | 77 | 778 | 200 | 71 |
| Chaliyar- PA | 744 | 84 | 65 | 492 | 186 | 73 |
| Chaliyar- FL | 321 | 61 | 21 | 178 | 121 | 13 |

PA- Particle-associated; FL- Free living

**Table 3.** Percentage abundance of different bacterioplankton in both Anjarakkandi and Chaliyar river, Kerala, India.

|  |  | Anjarakkandi | Chaliyar |
| --- | --- | --- | --- |
| Phylum | Proteobacteria | 67.07 | 46.35 |
|  | Firmicutes | 17.24 | 31.51 |
|  | Euryarchaeota | 6.12 | 10.68 |
|  | Chordata | 3.45 | 3.39 |
| Species | Marivivens sp. JLT3646 | 11.97 | 10.53 |
|  | Sulfitobacter sp. AM1-D1 | 11.97 | 3.2 |
|  | Bacillus subtilis | 16.19 | 10.07 |

## Discussion

Coastal waters, especially the estuarine regions are at a higher threat of eutrophication by various agricultural and other anthropogenic activities (Dai et al. 2017). Further, the report has also indicated that the bacterioplankton community structure is also changes in relation with the eutrophication status of the water body. The present study observed significantly higher number of proteobacteria, Actinobacteria, and Cyanobacteria in the estuarine waters of Anjarakkandi river, Dharmadam. Previous studies have indicated that the bacterioplankton belonging to the proteobacteria, Actinobacteria, and Cyanobacteria predominates in the eutrophic waters compared to other bacterial groups (Kiersztyn et al. 2019). The increased diversity of these bacteria correlates with the increased level of nitrogenous compounds present in the Anjarakkandi river and possibly an indicative of the eutrophication status of the estuary. Further, there observed significant elevation in the firmicutes diversity; these bacteria are present in the gut of humans and often indicates the faecal contamination in the water (Paruch et al. 2019a; Paruch et al. 2019b). Apart from that, there observed the presence of various nitrogen metabolizing bacteria including the roseobacter, Nitratiruptor and sulfitobacter. The Roseobacter is one the predominant bacterioplankton which is involved in the degradation various organic matter (Pohlner et al. 2019). Likewise, nitratiruptor is another nitrite reducing bacteria, which is active in the eutrophic waters (Nakagawa et al. 2005). It is thus possible that the increased occurrence of these bacterioplanktons in the Anjarakkandi river possibly indicate increased faecal contamination, subsequent increase in nitrogenous compounds and subsequent eutrophication.

**Figure 1.**
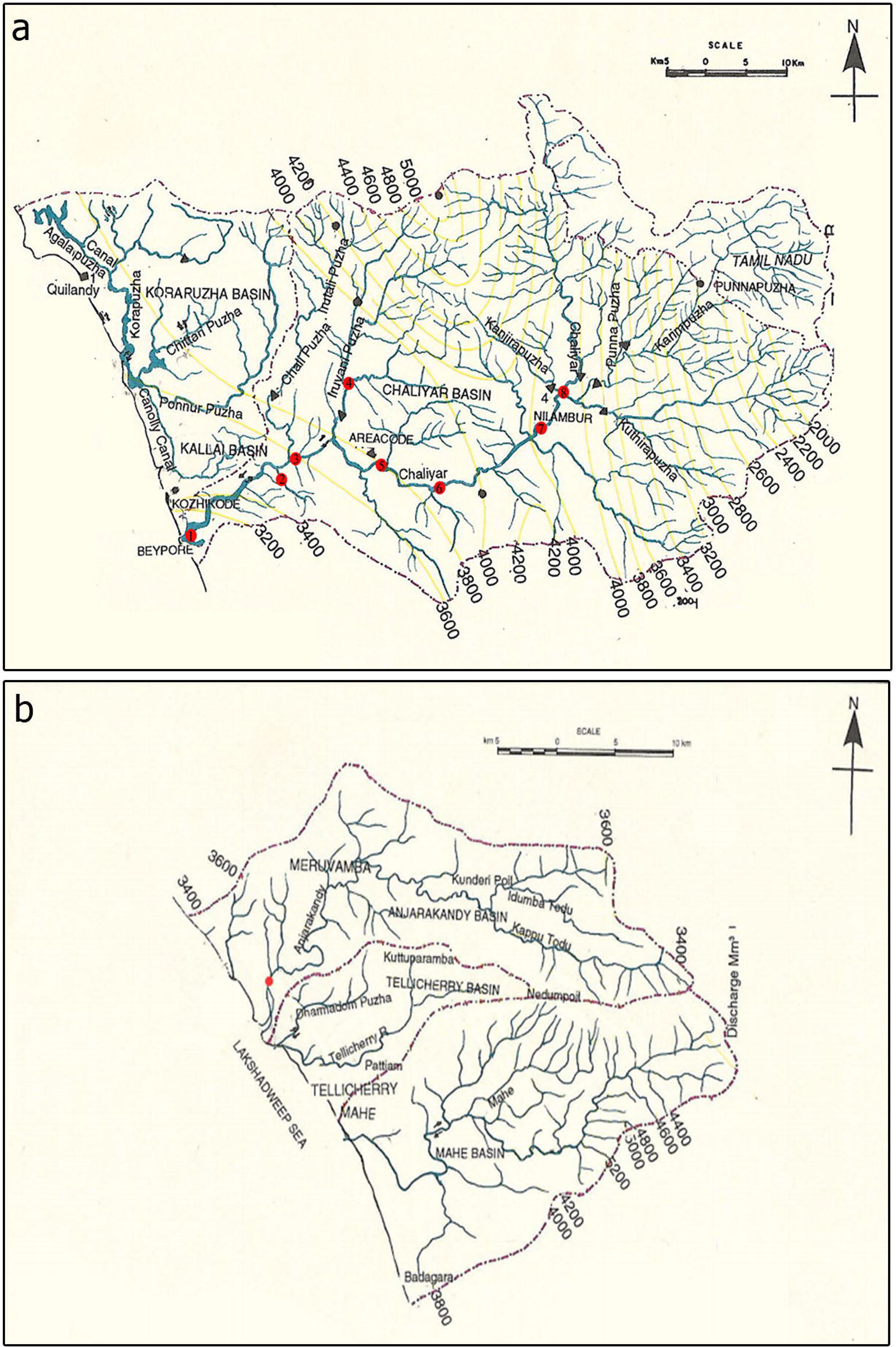
The site map of estuarine regions of Chaliyar river (a) and Anjarakkandi river (b).

## DECLARATIONS

### Availability of data and material

The data will be made available on valid request.

### Authors’ contributions

Jisha Jacob was contributed to the design, analysis of result, manuscript draft preparation, and statistical analysis. Nikhitha D conducted the experiments and carried out analysis and prepared the initial draft. Arunaksharan Narayanankutty contributed to the analysis of data and final manuscript editing. All authors have approved submission to the journal.

### Ethics approval

The study doesn’t involve any animal/ human subjects.

### Consent for publication

The study doesn’t use any third party material.

## Acknowledgement

The authors acknowledge Rashtriya Uchchathar Shiksha Abhiyan (RUSA) for financial support of the study. First author is thankful to the University grants commission for junior research fellowship.

