## Supplementary Material 1 for "Bacterioplankton diversity and pollution levels in the estuarine regions of Chaliyar and Anjarakkandi rivers, Kerala, India"

### Slide 1
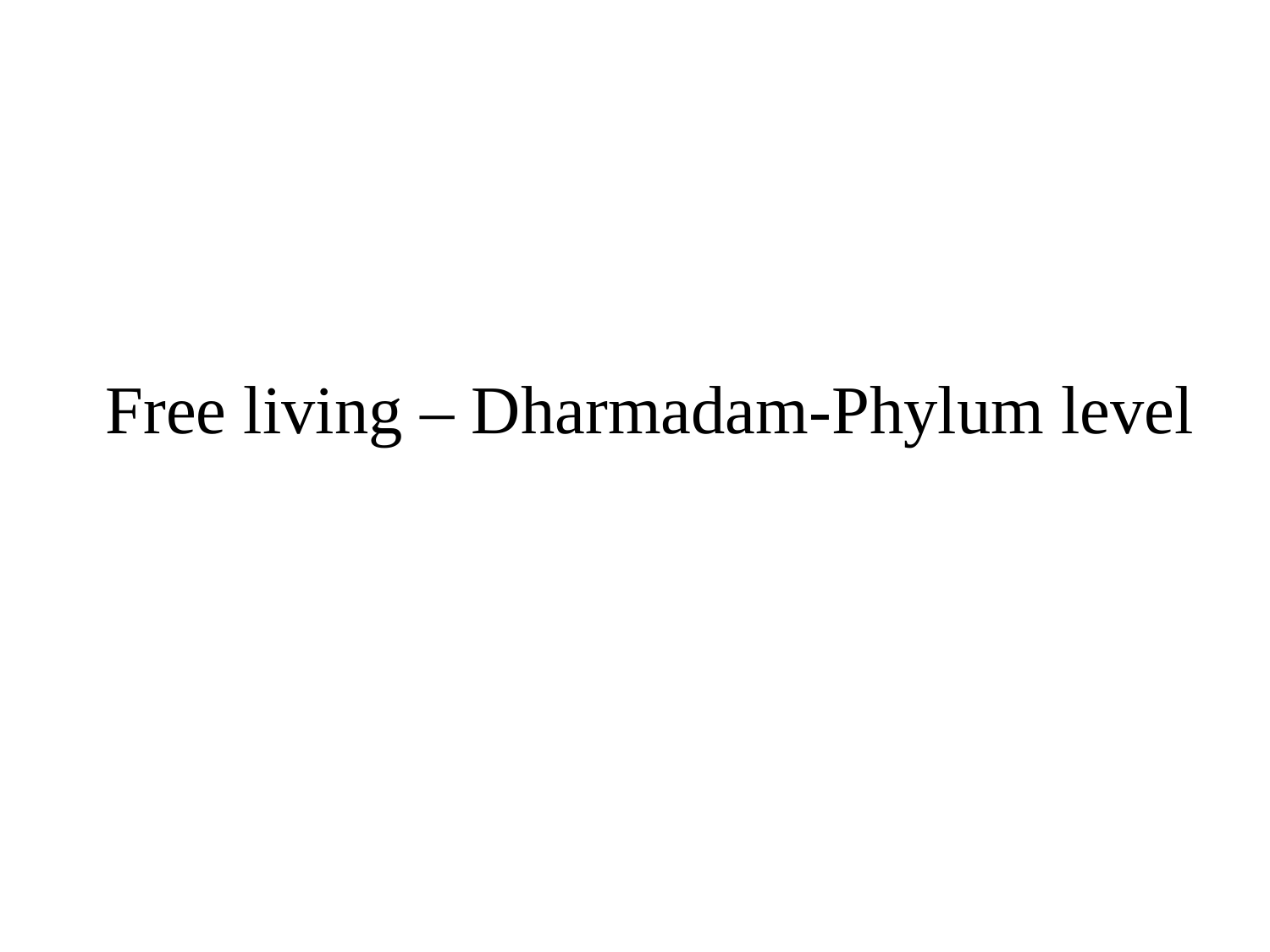

#### Free living – Dharmadam-Phylum level

### Slide 2
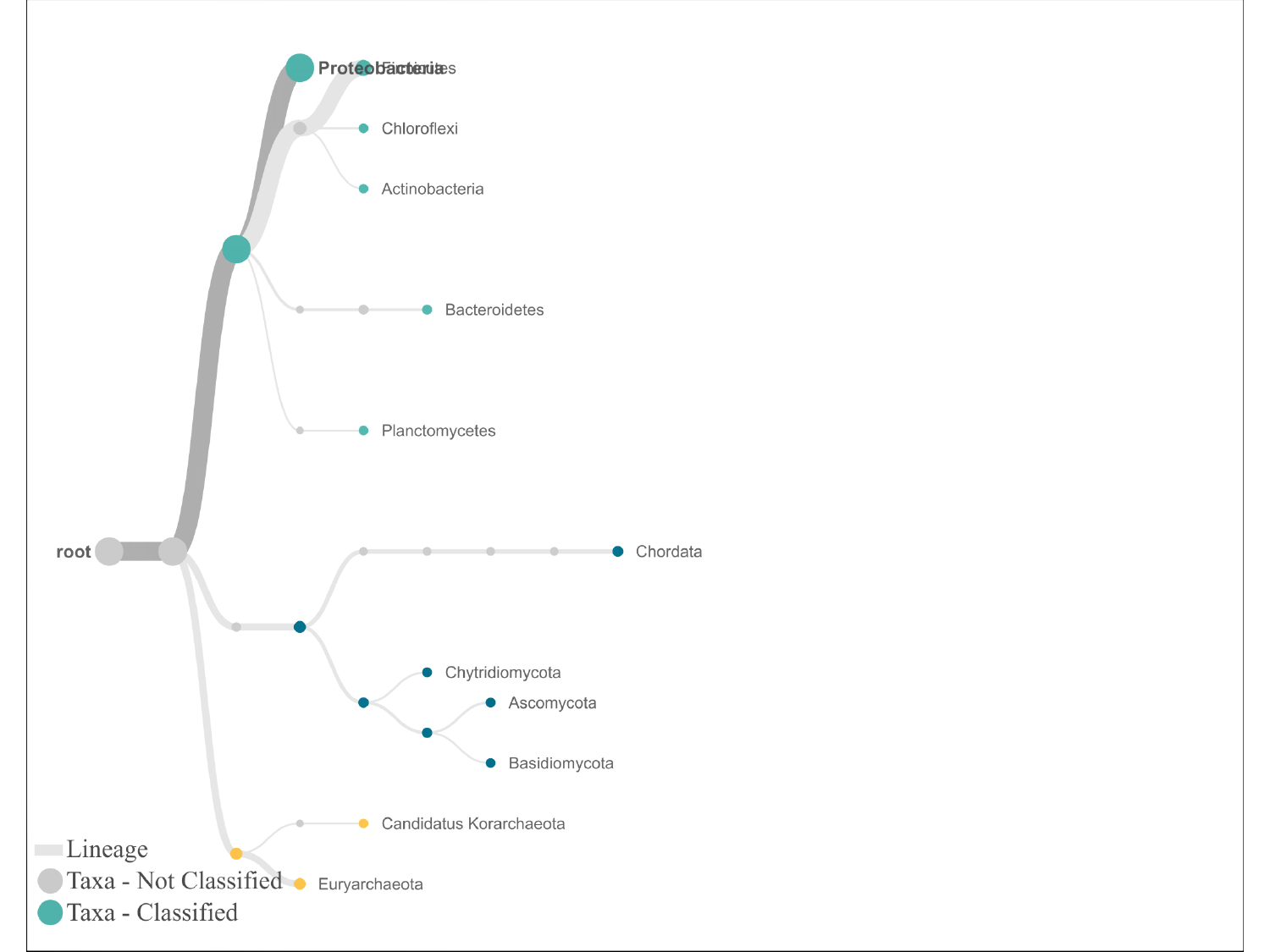

#

### Slide 3
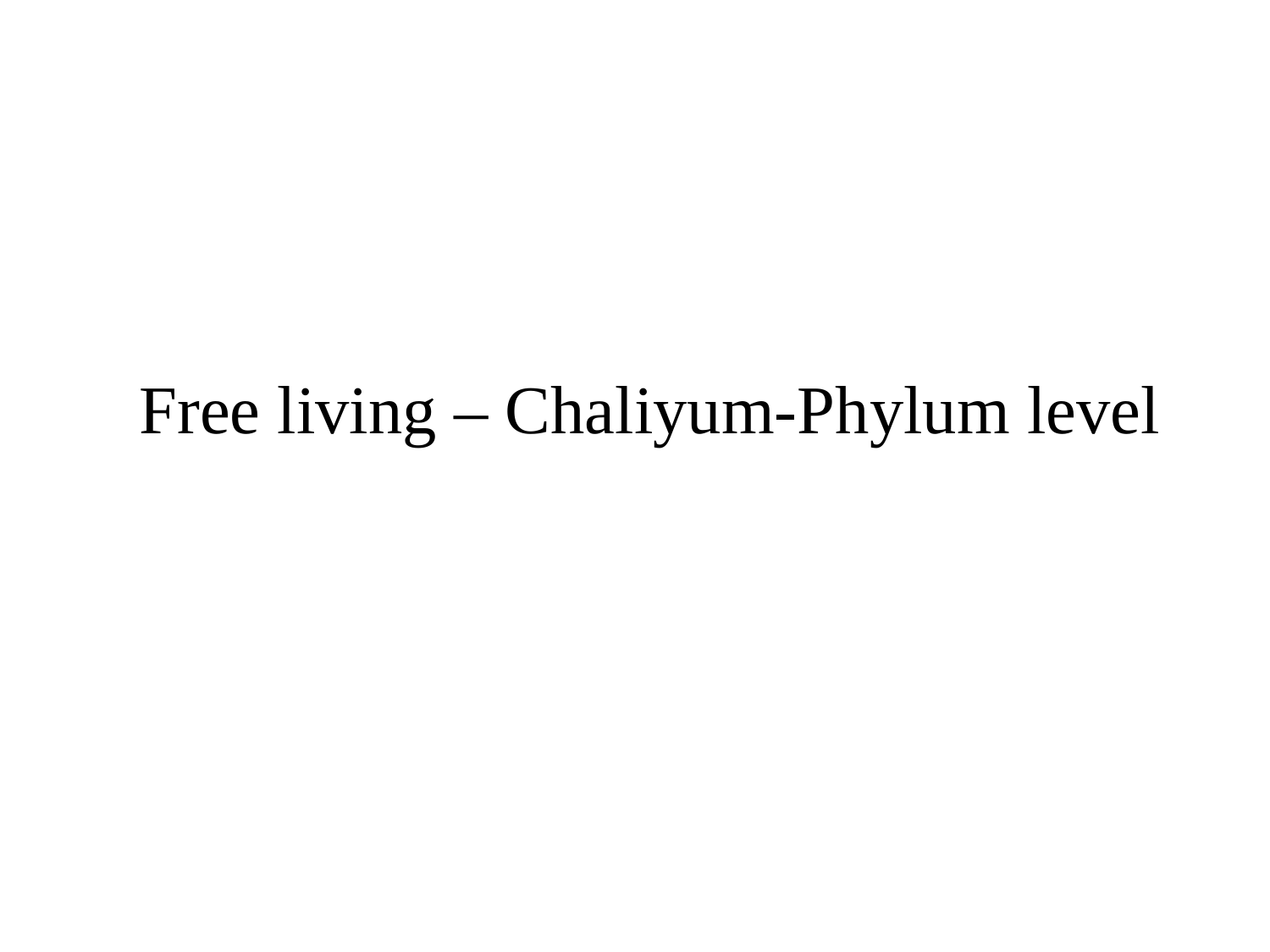

#### Free living – Chaliyum-Phylum level

### Slide 4
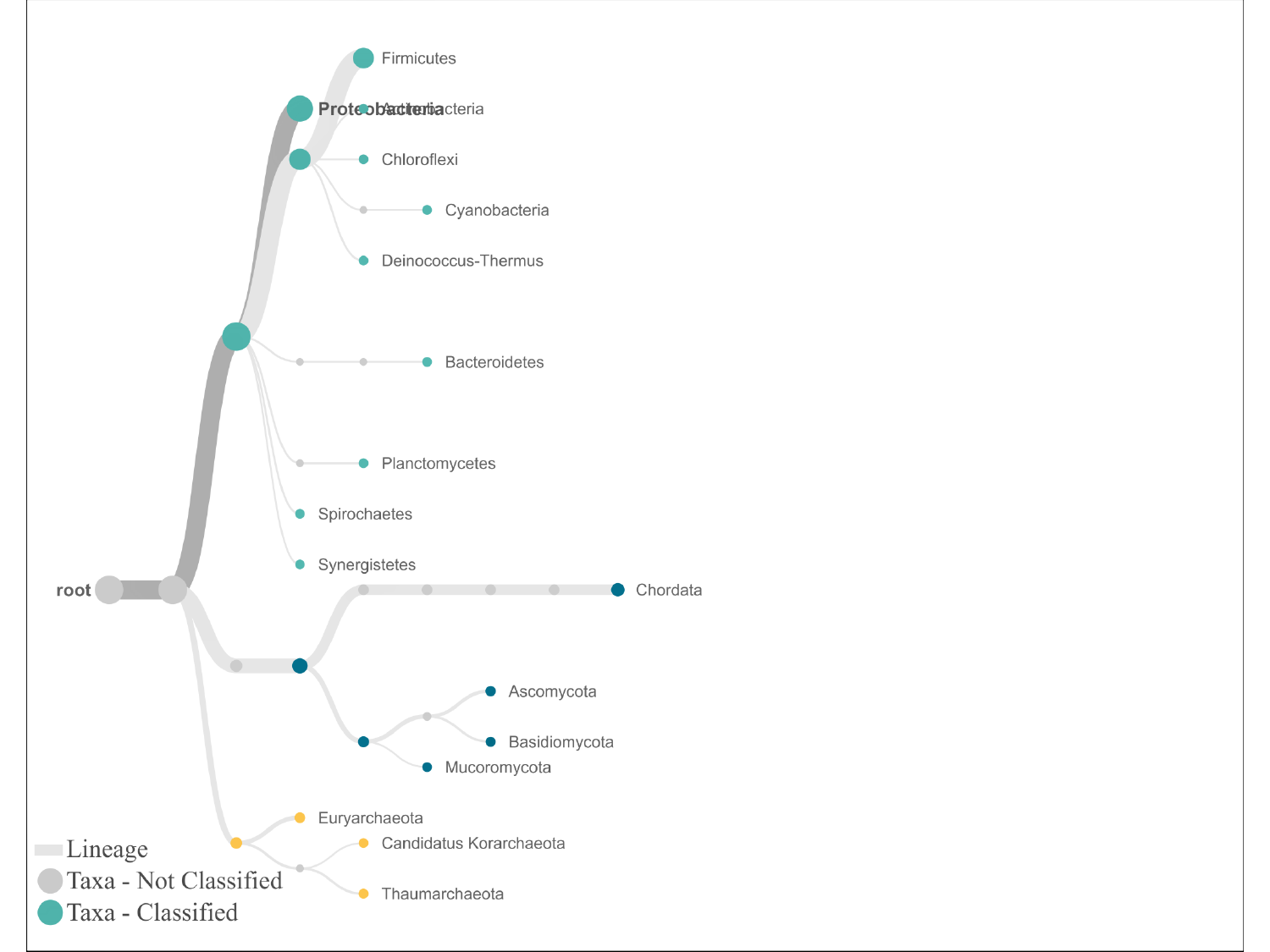

#

### Slide 5
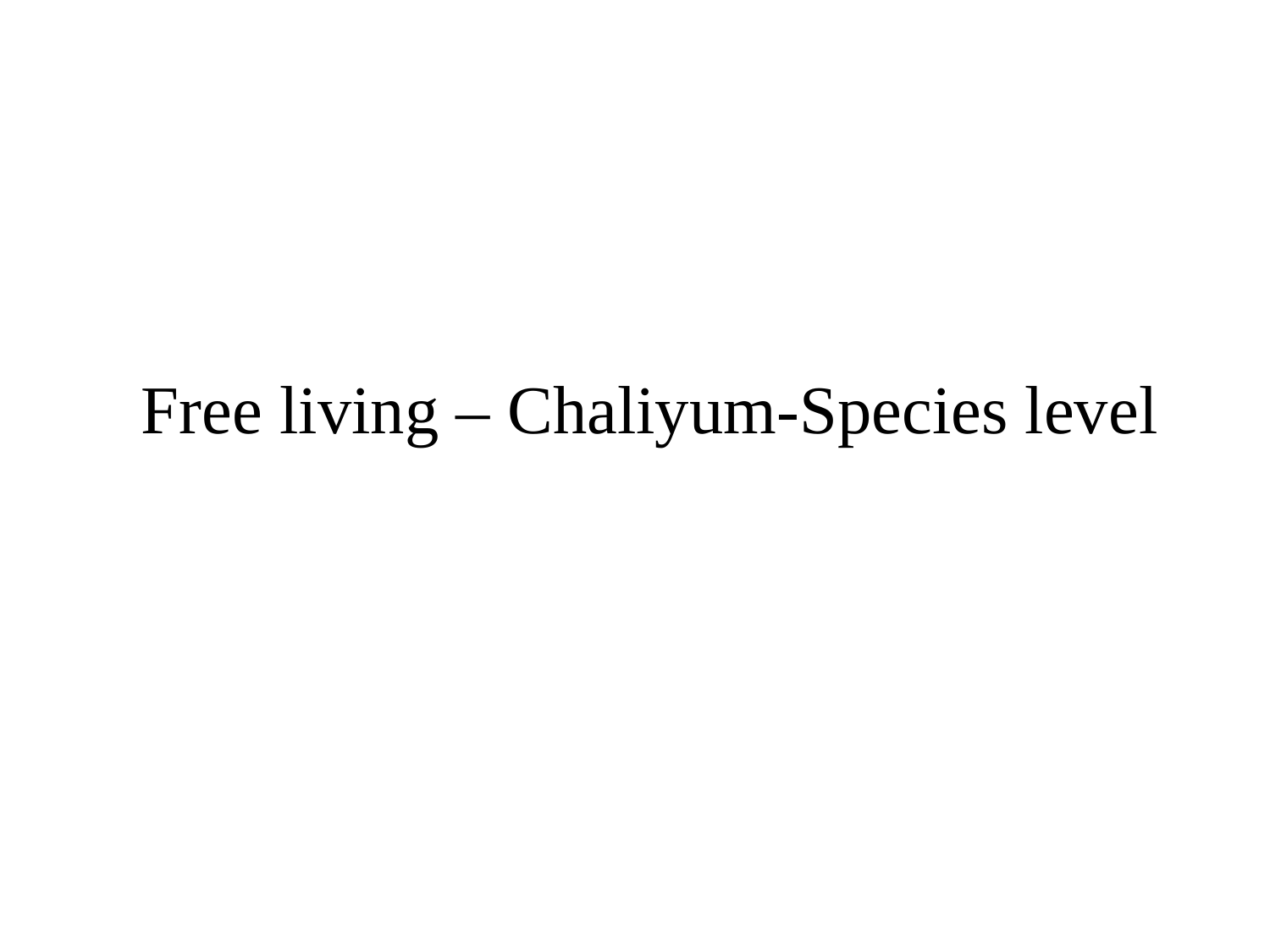

#### Free living – Chaliyum-Species level

### Slide 6
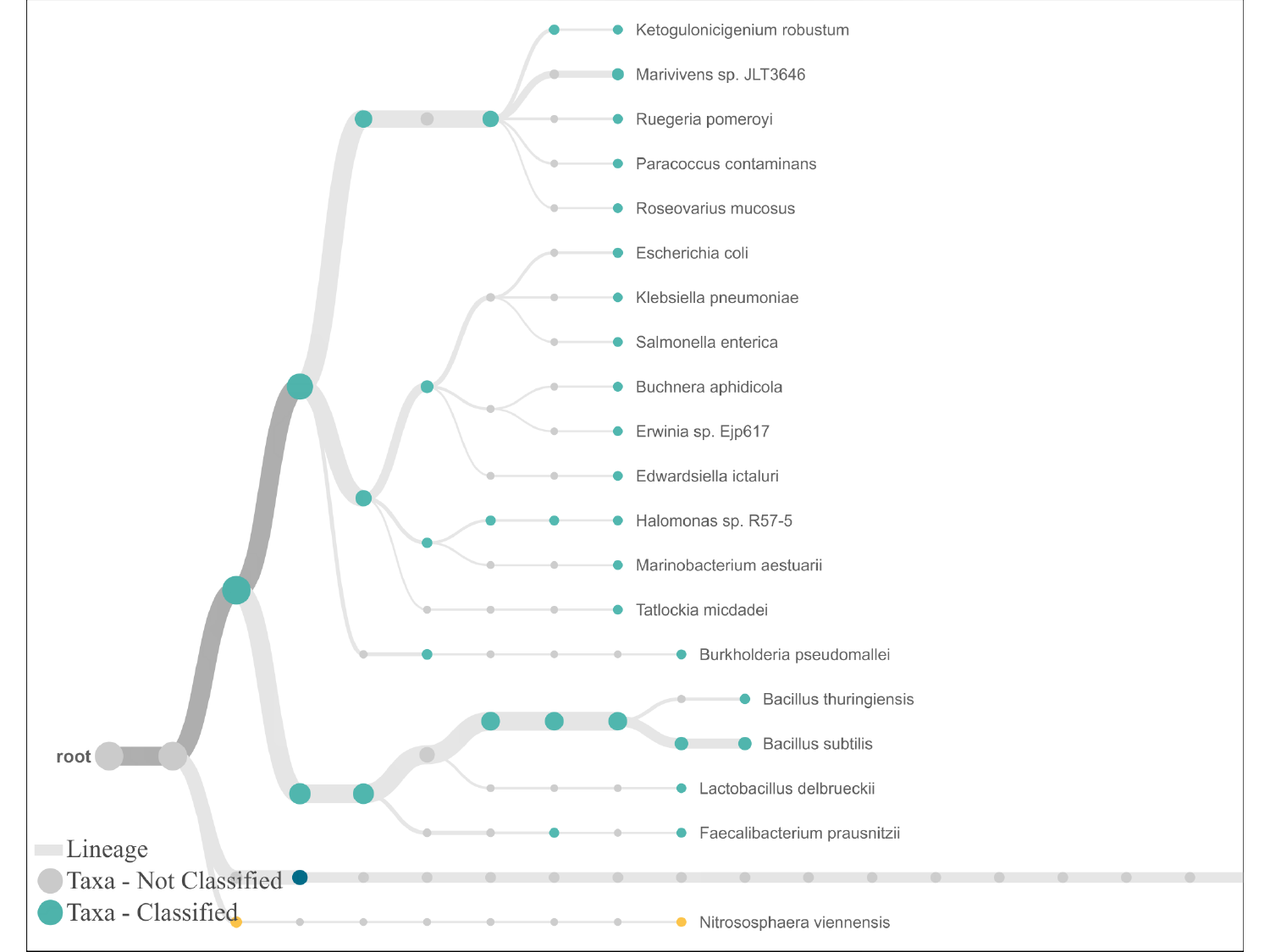

#

### Slide 7
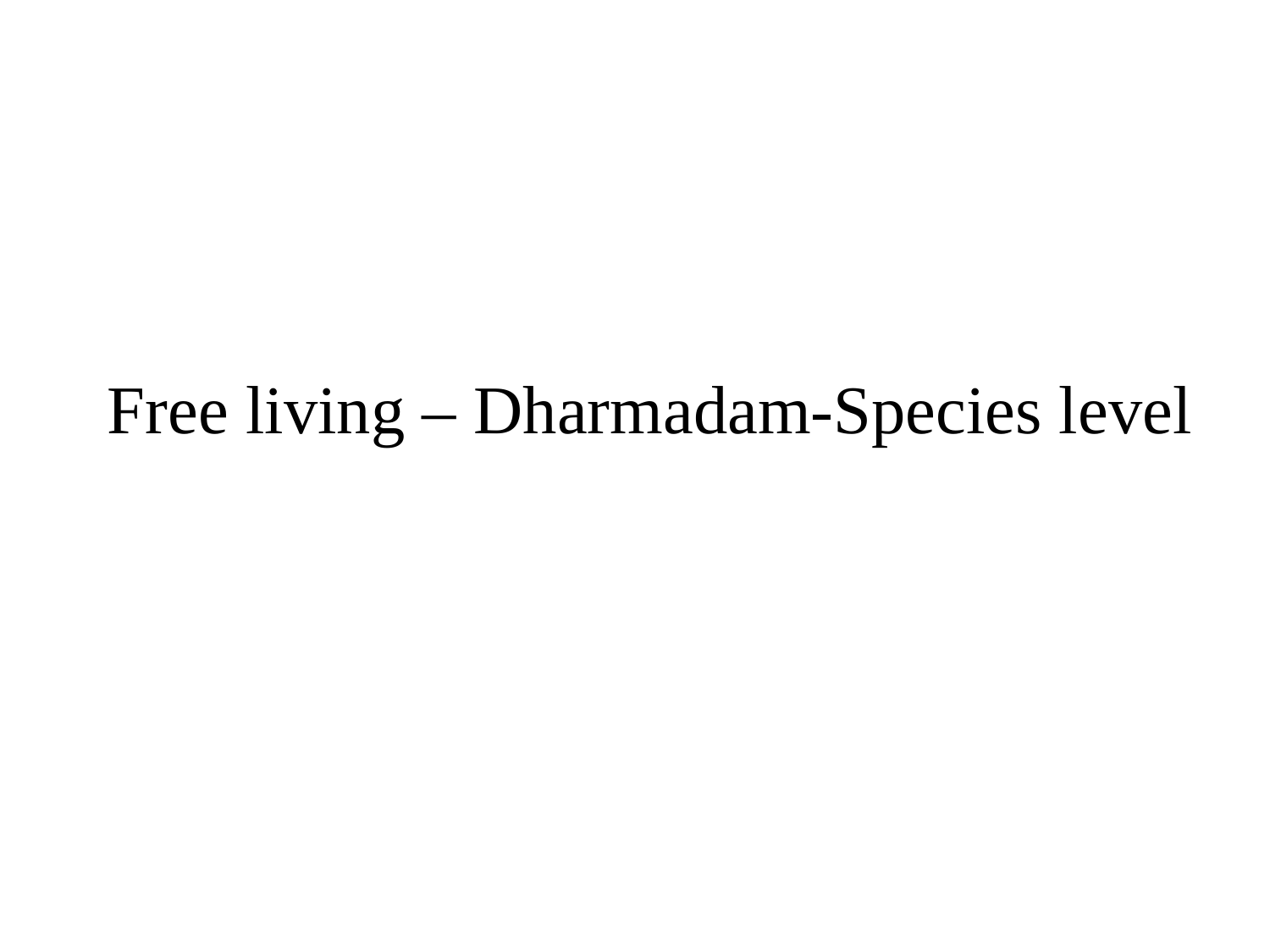

#### Free living – Dharmadam-Species level

### Slide 8
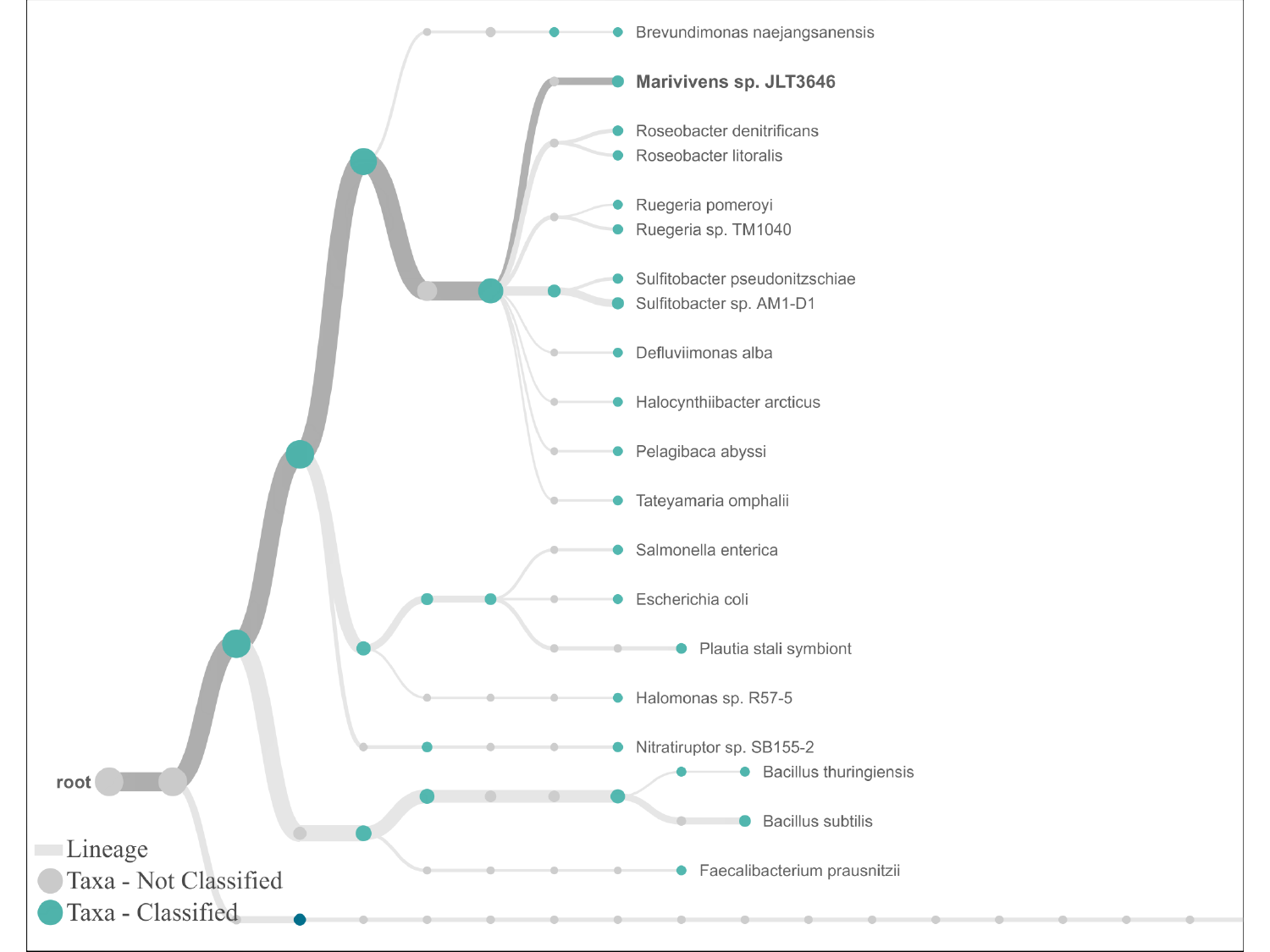

#

### Slide 9
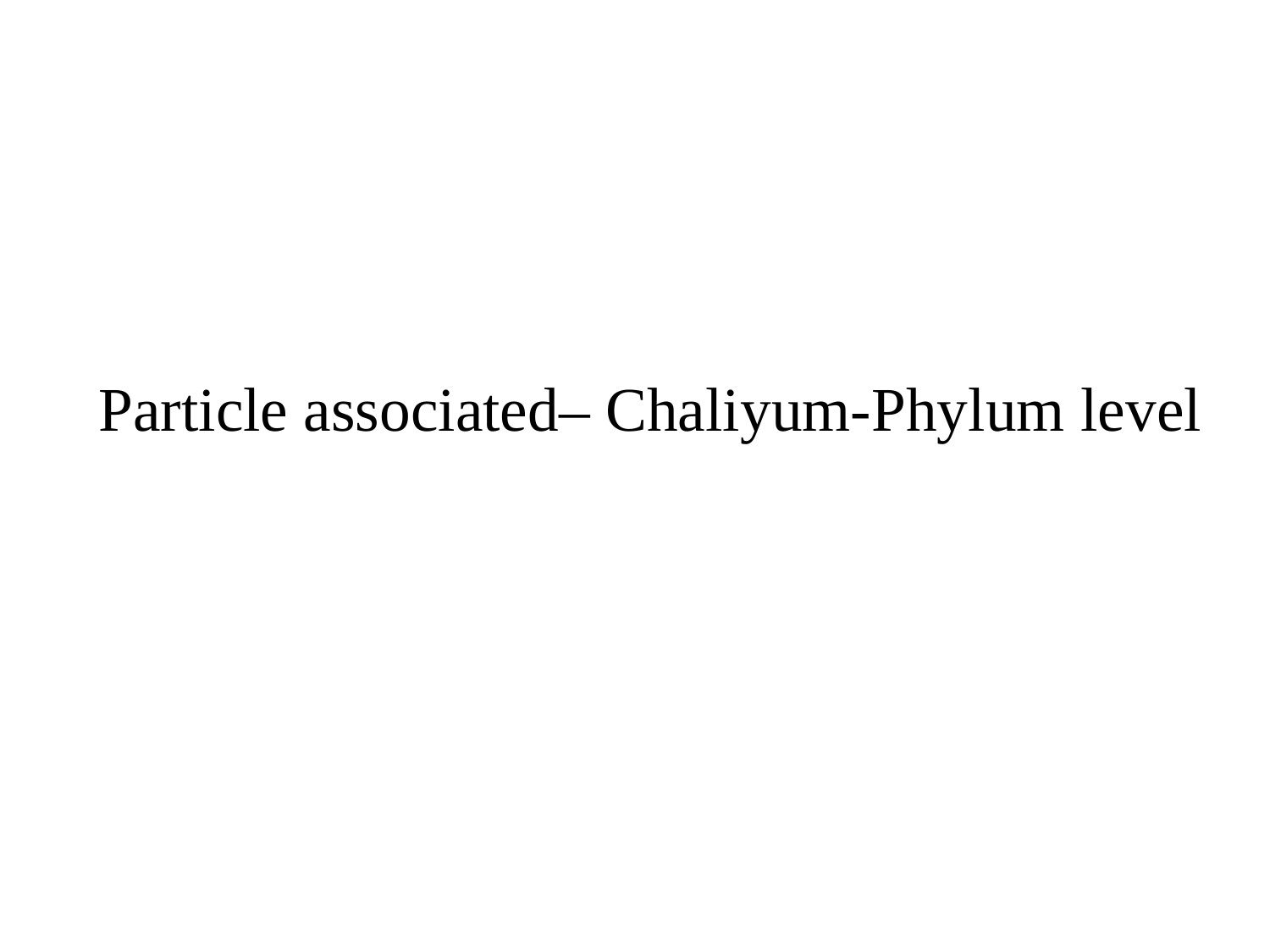

#### Particle associated– Chaliyum-Phylum level

### Slide 10
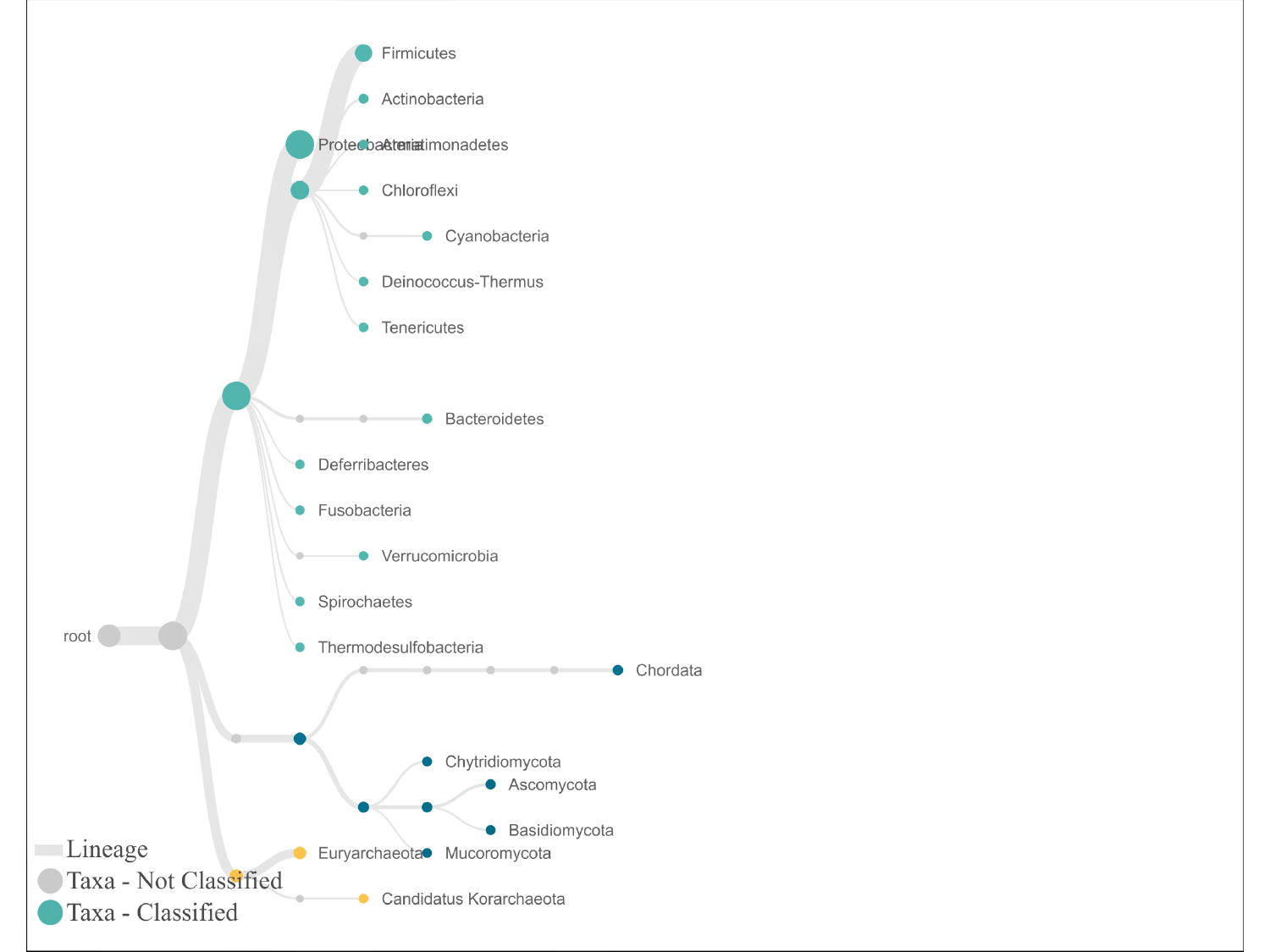

#

### Slide 11
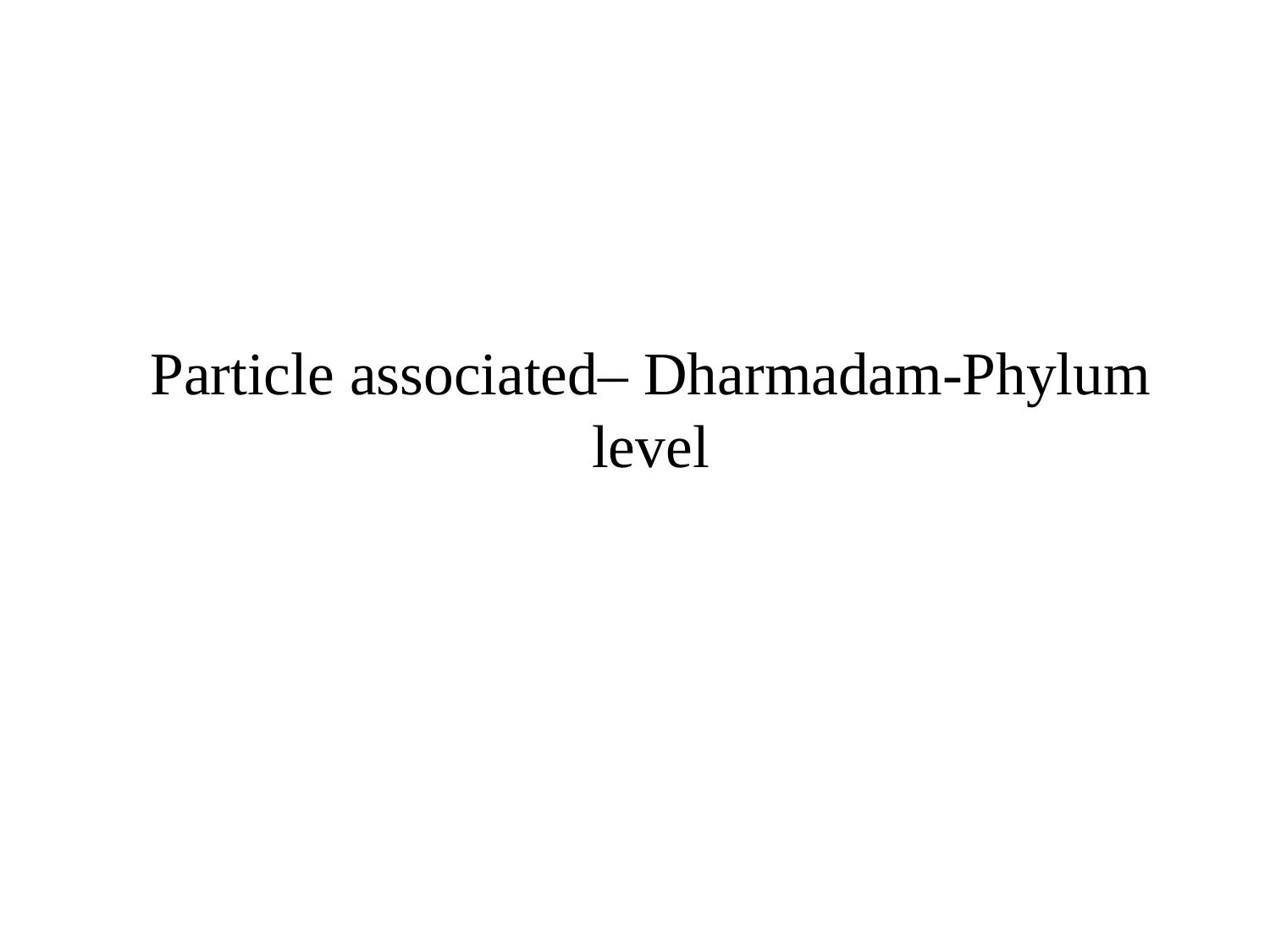

#### Particle associated– Dharmadam-Phylum level

### Slide 12
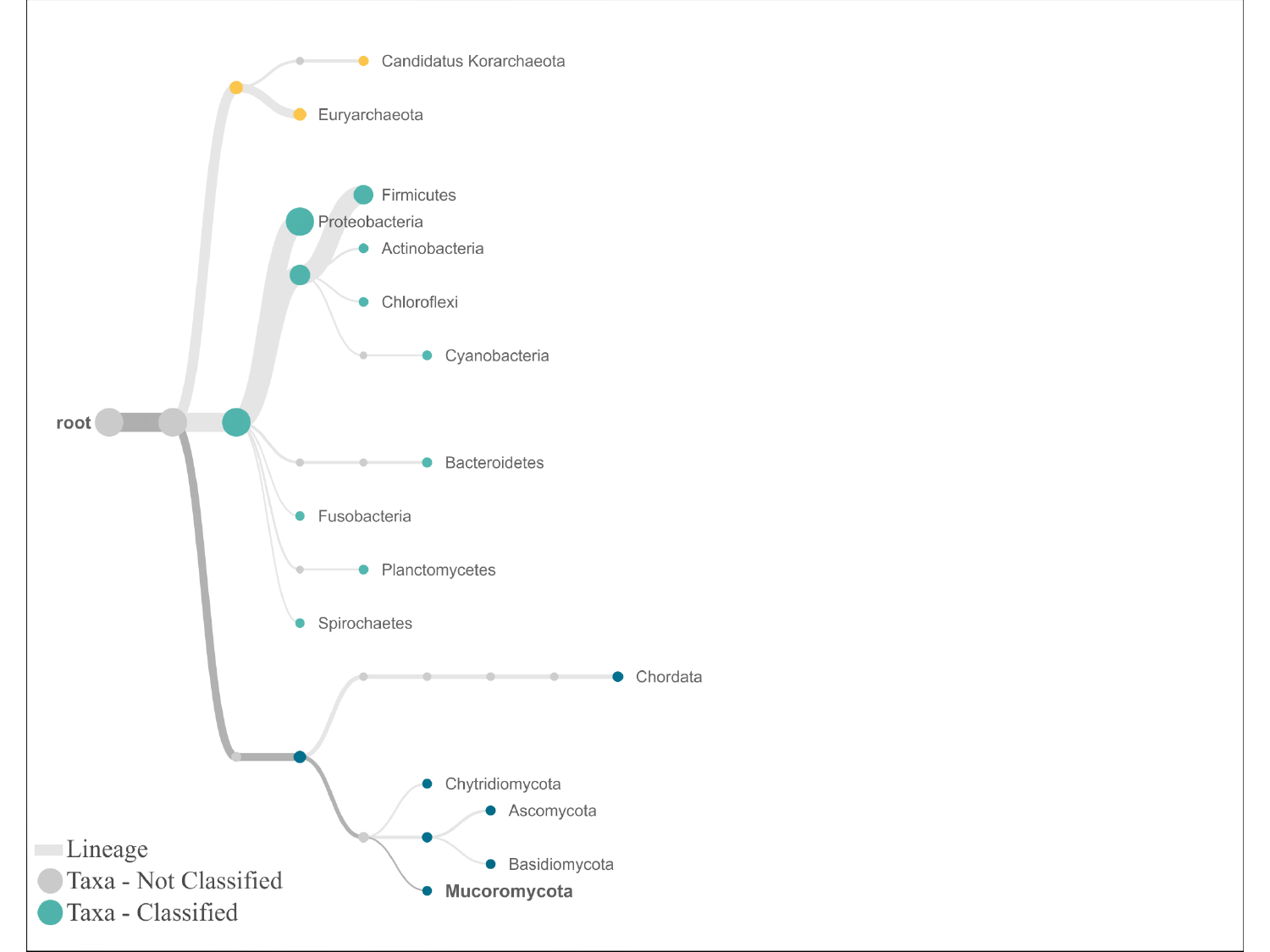

#

### Slide 13
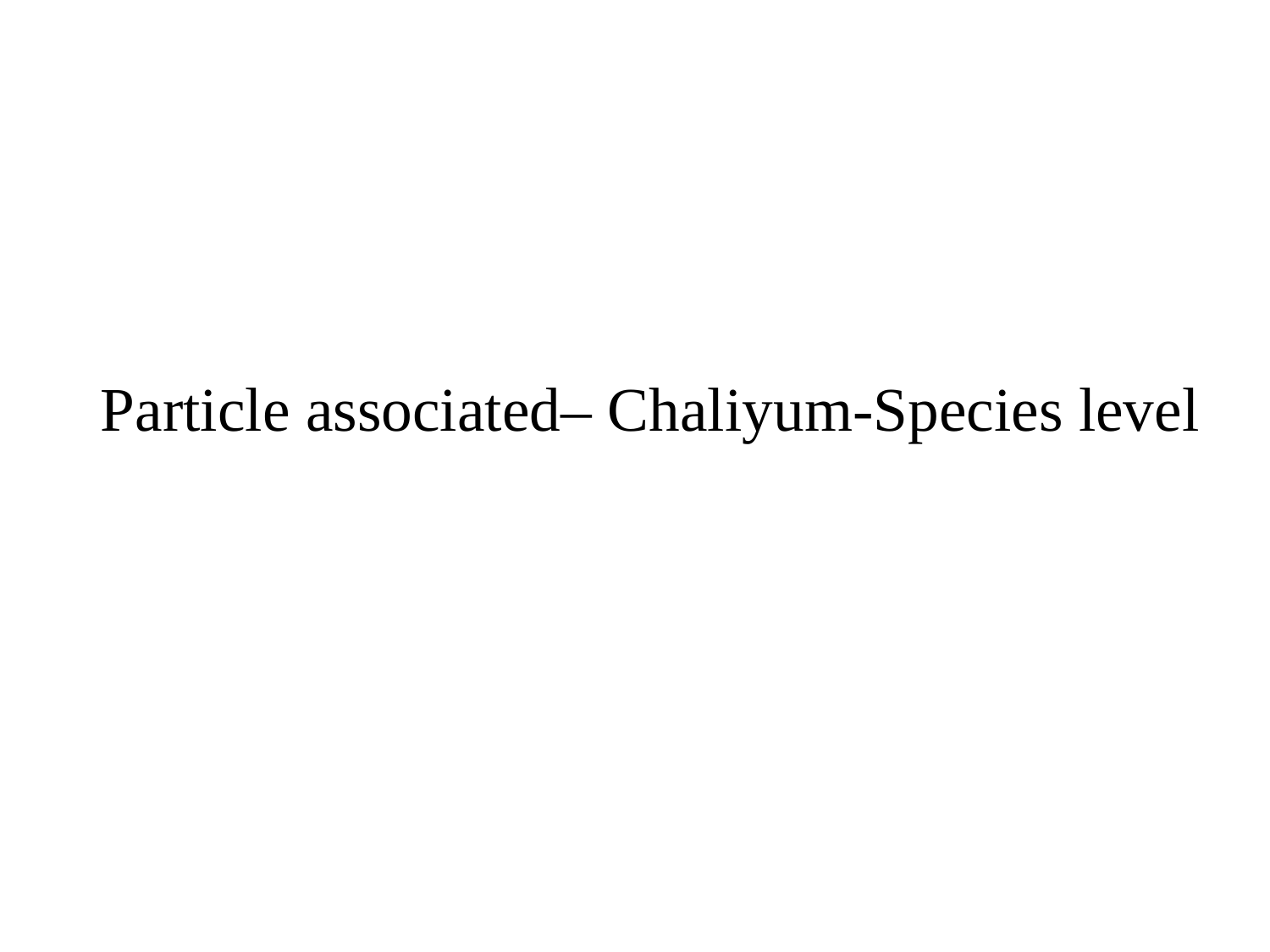

#### Particle associated– Chaliyum-Species level

### Slide 14
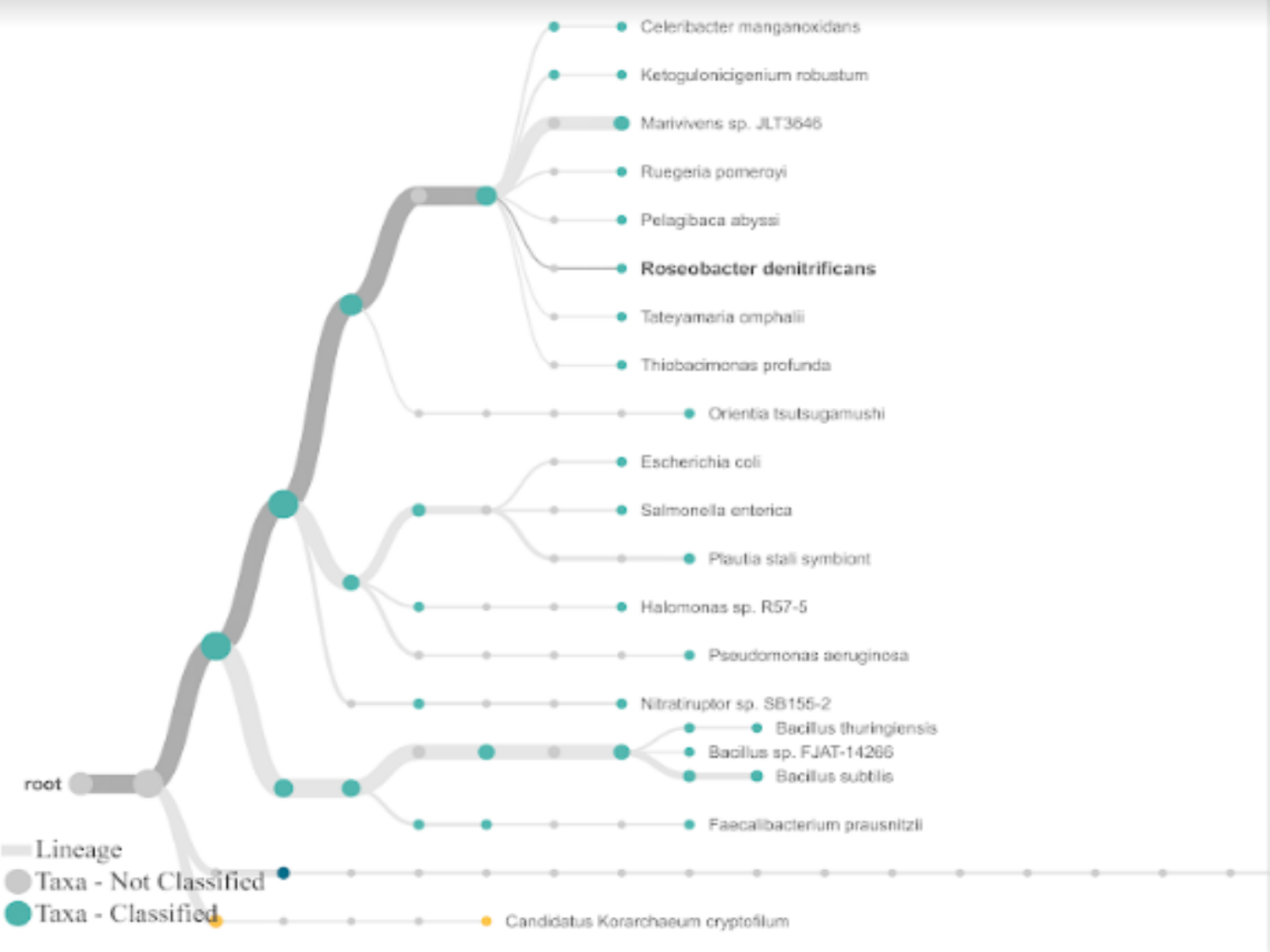

#

### Slide 15
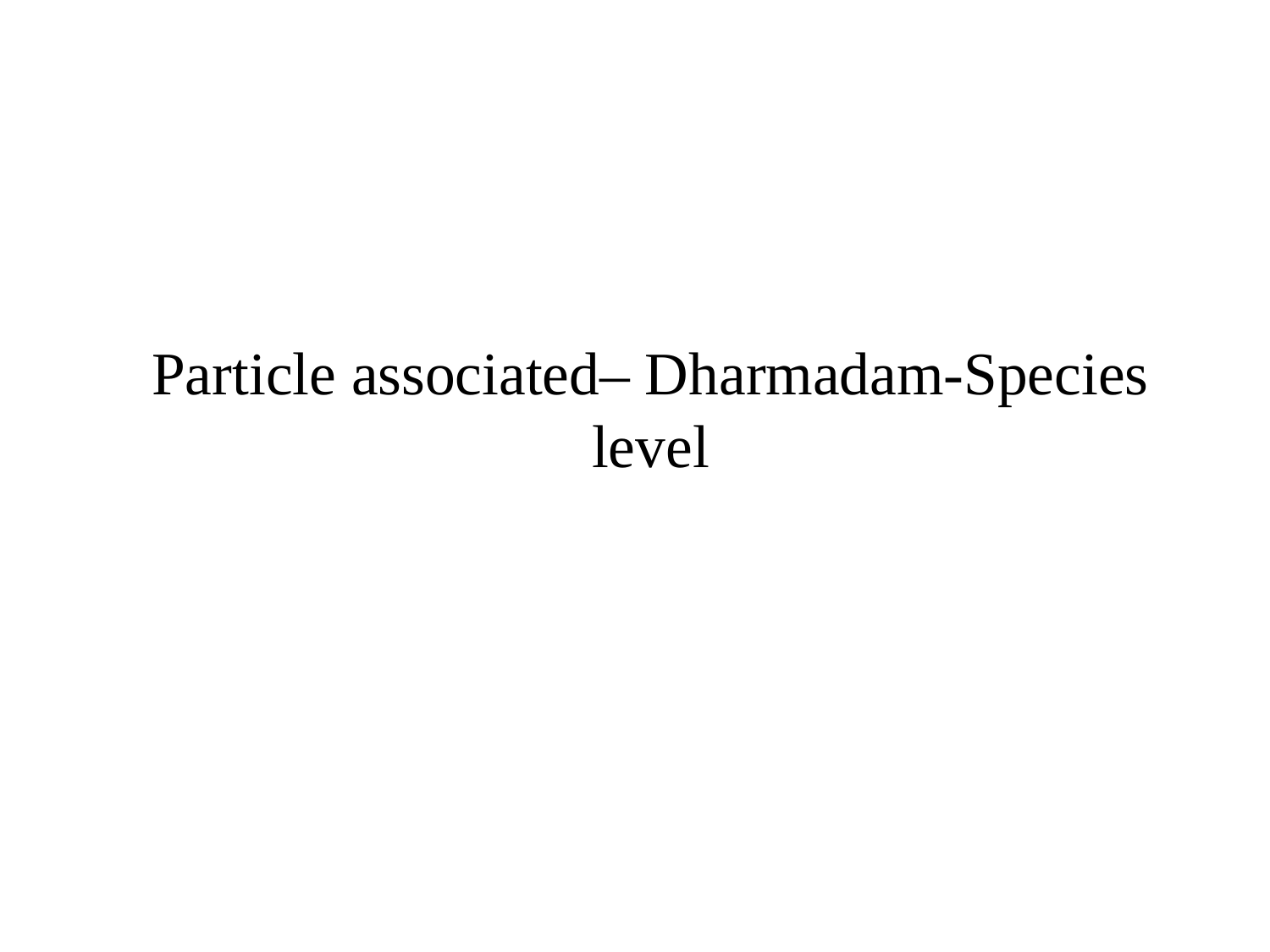

#### Particle associated– Dharmadam-Species level

### Slide 16
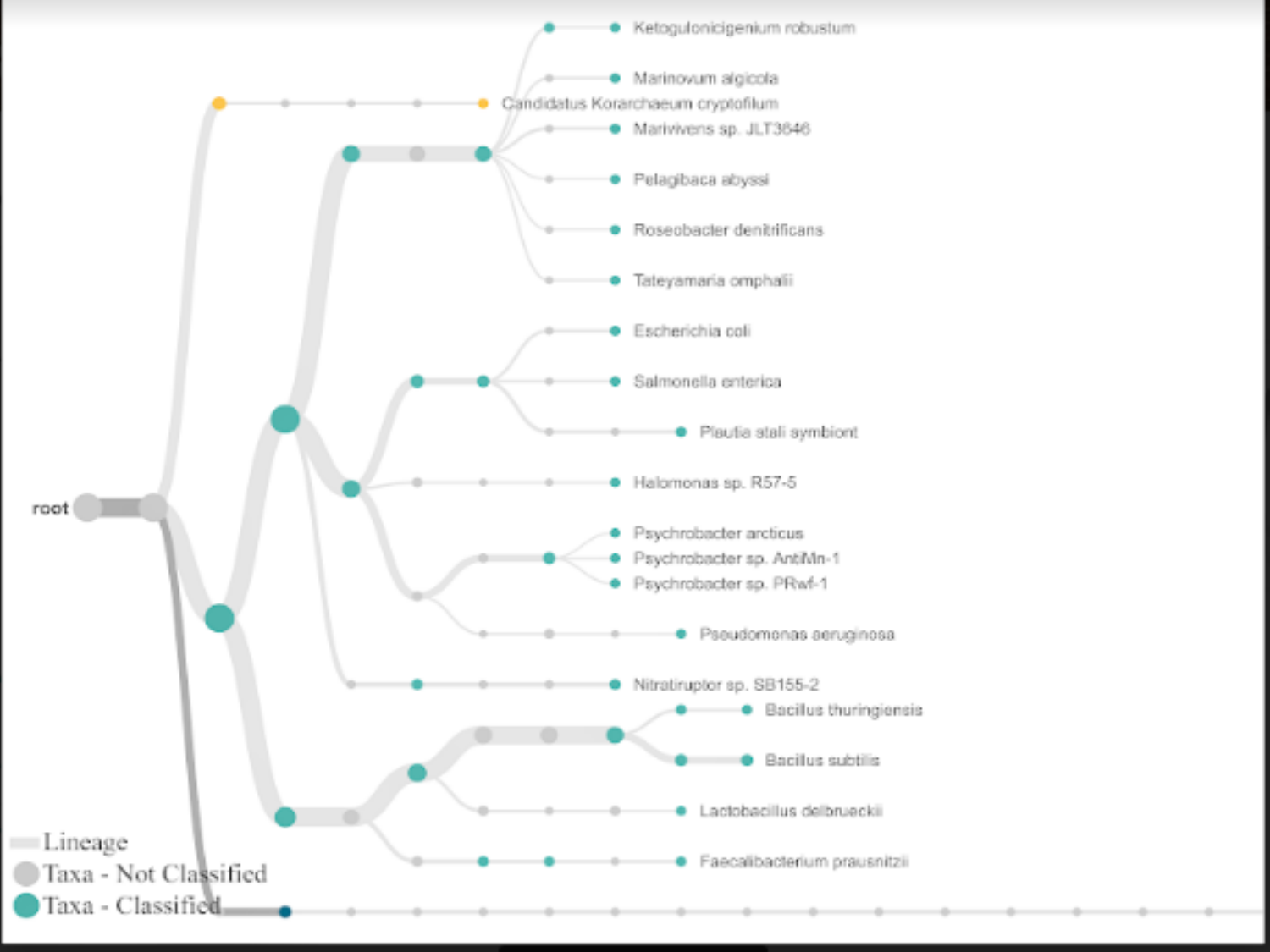

#
